# Tracing the biogeographical origin of South Asian populations using DNA SatNav

**DOI:** 10.1101/089466

**Authors:** Ranajit Das, Priyanka Upadhyai

## Abstract

The Indian subcontinent includes India, Bangladesh, Pakistan, Nepal, Bhutan, and Sri Lanka that collectively share common anthropological and cultural roots. Given the enigmatic population structure, complex history and genetic heterogeneity of populations from this region, their biogeographical origin and history remain a fascinating question. In this study we carried out an in-depth genetic comparison of the five South Asian populations available in the 1000 Genomes Project, namely Gujarati Indians from Houston, Texas (GIH), Punjabis from Lahore (PJL), Indian Telugus from UK (ITU), Sri Lankan Tamils from UK (STU) and Bengalis from Bangladesh (BEB), tracing their putative biogeographical origin using a DNA SatNav algorithm - Geographical Population Structure (GPS). GPS positioned >70% of GIH and PJL genomes in North India and >80% of ITU and STU samples in South India. All South Asian genomes appeared to be assigned with reasonable accuracy, along trade routes that thrived in the ancient Mauryan Empire, which had played a significant role in unifying the Indian subcontinent and in the process brought the ancient North and South Indian populations in close proximity, promoting admixture between them, ~2300 years before present (YBP). Our findings suggest that the genetic admixture between ancient North and South Indian populations likely first occurred along the Godavari and Krishna river basin in Central-South India. Finally our biogeographical analyses provide critical insights into the population history and sociocultural forces driving migration patterns that may have been instrumental in shaping the population structure of the Indian subcontinent.

## Introduction

India is the seventh largest country in the world in terms of land area; it lies in the Indian subcontinent that also includes its neighboring countries, Bangladesh, Pakistan, Nepal, Bhutan and Sri Lanka, which collectively share common anthropological and cultural roots. India alone is the second most populous country in the world with more than 4000 anthropologically distinct groups, 22 languages and several other regional dialects.^1,2^ The Indian subcontinent is unique in its complex history, diversity and eclectic population structure, thus, the geographic origins of populations from this region remain a challenging question in genetics, cultural anthropology and history.

The complex tapestry of genetic and cultural diversity observed in contemporary India is a result of admixture between its autochthonous and immigrant gene-pools. Several lines of evidence have suggested that by 50,000 to 20,000 years before present (YBP) modern humans had migrated to various parts of India and its surrounding areas in South Asia.^3,4^ Linguistic and genetic evidences suggest that this was followed by the early migration of putative proto-Dravidian language speakers, ~ 6000 YBP from West Asia (Elam province), eventually giving rise to the Dravidian speaking, South Indian populations.^5-7^ The ancient peopling of India was followed by several more recent waves of immigration from West and East Eurasia (Central Asia, the Middle East, Europe and the Caucasus) by populations that spoke Indo-European languages and who are often presumed to be ‘Aryans’.^8-11^ The migration of Eurasian populations into the Indian sub-continent has often been ascribed to purported Aryan invasions and/or the spread of agriculture.^12-15^ The later Eurasian immigrants diffused throughout the Indian subcontinent and are believed to have either admixed with the existing Indian populations, primarily contributing to the North Indian gene-pool.^7,16^ Several lines of evidence based on analyses of Y chromosome, mitochondrial DNA^9,17-19^ and whole-genome approaches^10,20-22^ have indicated that the modern populations of India are descendants of two genetically divergent ancestral populations, the Indo-European language speaking Ancestral North Indians (ANI) and the Dravidian speaking Ancestral South Indians (ASI).

Genome-wide analyses of individuals from well-defined ethno-linguistic groups of South Asia estimated that the admixture between ancestral North and South Indian populations likely occurred ~1900 to 4200 YBP.^23^ Despite the high admixture between ancestral Indian populations, the ensuing sociocultural stratification resulted in more than 400 tribal and 4000 caste groups, that are endogamous, in present day India.^24^ Strictly enforced sociocultural norms barred mate-exchange between individuals belonging to distinct strata of the society and resulted in consanguinity being as high as 20-30% in some Indian populations, leading to the possibility of founder mutations and a high prevalence of recessive inherited diseases in these groups.^22,25^ Thus, population stratification and constrained gene-flow between distinct groups altered the demography and genetic structure of populations from the Indian subcontinent.

Several studies have used high-resolution genome-wide approaches to assess the genetic heterogeneity and diversity of South Asian populations. The Indian Genome variation consortium (2008) analyzed 405 single nucleotide polymorphisms (SNPs) from 75 genes in 55 endogamous Indian populations and observed high levels of genetic divergence between them based on ethnicity and language.^2^ Reich *et al*. (2009) investigated the complex genetic canvas of India using a much larger sample size, wherein 560,123 SNPs were surveyed in 132 Indians originating from 25 ethno-linguistically and socially distinctive groups across the country.^22^ Another study compared 85 Gujaratis or North Indian genomes from Phase 3 of the International HapMap project to 83 South Asian Indians from Singapore to determine recent positive selection on the *SLC24A5* gene that plays a role in skin pigmentation, suggesting a higher degree of genetic closeness between Gujaratis and Europeans^26^.

In the present study we have carried out an in-depth genetic comparison of the five South Asian populations available in 1000 Genomes project, namely Gujarati Indians from Houston, Texas, USA (GIH), Punjabis from Lahore, Pakistan (PJL), Indian Telugu from UK (ITU), Sri Lankan Tamil from UK (STU), and Bengalis from Bangladesh (BEB), assessing a total of 489 (103 GIH, 96 PJL, 102 ITU, 102 STU and 86 BEB) samples.^27^ We also investigated several previously reported Indian populations from distinct linguistic groups (N = 378), obtained from the Reich lab at Harvard Medical School, USA.^23^ Overall we evaluated 494,863 previously assessed SNPs^23^ to interrogate the genetic diversity among the above mentioned five South Asian populations. Given the esoteric population history and profound genetic diversity in the Indian subcontinent, questions regarding their origin remain particularly fascinating. To facilitate our understanding of the biogeographic origin of South Asian populations, we employed an admixture-based method, known as Geographical Population Structure (GPS)^28^, which has been successfully applied to trace the accurate origin of various modern populations including Yiddish language speakers^29^ and the Mountain Druze population^30^.

## Materials and Methods

### Datasets

The dataset used in the present study was obtained from 1000 Genomes project phase 3 and from the Reich Lab, Department of Genetics, Harvard Medical School, USA.^21,23^ The original data obtained from the Reich lab was in the eigenstrat format and was converted into the PLINK bed format using EIG v4.2.^31^ The 1000 Genomes dataset was in the VCF format and was converted to PLINK format using VCF tools.^32^ Thereafter data pertaining to all South Asian genomes (N = 489) were retained using PLINK ‐‐keep command and 494,863 SNPs previously assessed^23^ were extracted using PLINK ‐‐extract command (http://pngu.mgh.harvard.edu/purcell/plink/)^33^ and subsequently converted into a bed file. Before merging the 1000 Genomes data file with that obtained of the Reich lab, DNA strands were flipped for 241,785 SNPS in the Reich Lab dataset using PLINK ‐‐flip command to ensure that the two datasets were compatible with each other. Finally, the bed files from 1000 Genomes project and the Reich Lab were merged using PLINK ‐‐ bmerge command.

### Population clustering and Admixture analysis

Population stratification was estimated using the ‐‐cluster command in PLINK. Multidimensional scaling analysis was performed in PLINK using ‐‐mds command alongside the ‐‐read-genome flag. The multidimensional (mds) plot was generated in R v3.2.3 comprising of the 489 South Asian samples from 1000 Genomes project. The ancestry of all 489 samples derived from the 1000 Genomes project and 378 samples from the Reich Lab dataset was estimated using ADMIXTURE v1.3.^34^ The number of ancestral populations (K) was estimated using the cross-validation procedure implemented in ADMIXTURE v1.3 using a ‐‐cv flag to the ADMIXTURE command line. The lowest cv error was estimated for K = 5.

### Tracing the biogeographical origin of South Asian populations using GPS algorithm

Biogeographical analysis was performed using the Geographic Population Structure (GPS) algorithm that has been shown to be highly accurate when compared to alternative approaches like spatial ancestry analysis for biogeography and has been demonstrated to effectively trace the origins of several modern populations^29,30,35,36^ Given a sample of unknown geographic origin and admixture proportions that correspond to putative ancestral populations, the GPS tool converts the genetic distances between the test sample and the nearest ancient reference populations into geographic distances. The coordinates predicted through GPS should be considered as the last place where the admixture had occurred or the ‘geographical origin’. All supervised admixture proportions were calculated as in Elhaik *et al.* (2014).^28^ Essentially, the GPS algorithm correlates the admixture patterns of individuals of unknown origin with geographical coordinates using the admixture fractions (GEN file) and geographical locations (GEO file) of reference samples. In the present study we used the Indian populations described in Moorjani *et al.*, 2013^23^ as the reference and interpreted their admixture fractions and geographic locations (latitudnal and longitudnal coordinates) to determine the biogeographical ancestry of five South Asian populations (N = 489) derived from the 1000 Genomes project. Thus, the GEN file contains five admixture coefficients corresponding to 378 individuals across 52 reference populations from around India and the GEO file contains the geographic coordinates (latitude and longitude) for the same individuals.

### Determining the accuracy of GPS prediction

Geographic distances (‘Laws of cosines’, great circle distance) between the physical location of the 1000 Genomes samples and their corresponding GPS assigned locations was calculated using the R package geosphere (https://CRAN.R-project.org/package=geosphere). Since GIH, ITU, and STU genomes were of the South Asian diaspora residing outside the Indian subcontinent, the capital cities of their ancestral region or country, Ahmedabad, Hyderabad, and Colombo respectively were used as their native regional location. For PJL and BEB samples the geographic coordinates of Lahore and Dhaka, respectively were used for the geographic distance calculations.

## Results

### Clustering of populations

The multidimensional plot revealed three distinct clusters: an isolated cluster of Gujaratis (GIH) of North Indian descent, a small cluster comprising of Telugus (ITU) of South Indian descent, and a large cluster that included individuals of all remaining populations (Figure 1). Bengali (BEB) samples largely overlapped with Telugu (ITU) and Tamil (STU) samples. Punjabis (PJL) showed the highest genomic versatility, overlapping with both Gujaratis as well as the Tamil-Telugu-Bengali cluster. The Gujaratis showed an interesting clustering pattern, wherein 57 GIH samples formed a distinct homogeneous cluster while the rest overlapped with the larger South Asian cluster. Likewise, 15 Telugu genomes also formed a homogeneous cluster distinct from the South Asian milieu while the rest overlapped with the latter.

**Figure 1.**
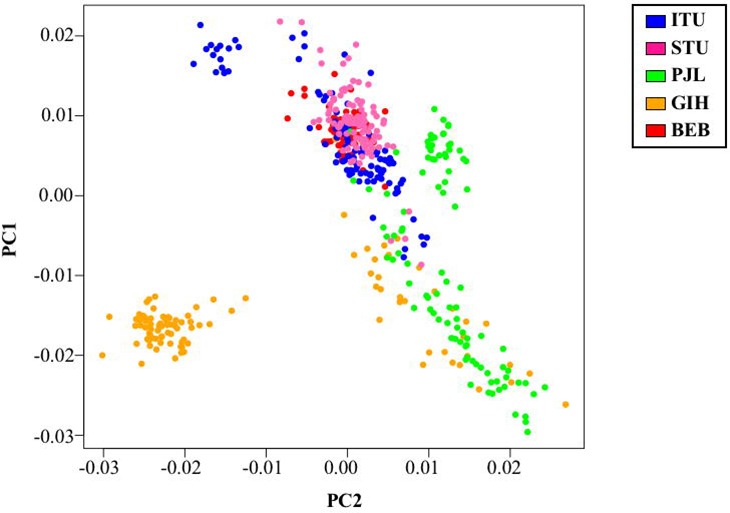
A multidimensional scaling plot of five South Asian populations from 1000 Genomes project. In this scatter plot each point represents an individual. Multidimensional scaling analysis was performed in PLINK and the plot was generated in R v3.2.3. The red, blue, orange, green, and pink circles designate BEB, ITU, GIH, PJL, STU populations respectively.

At K=5, a discernible degree of genetic admixture between the North and South Indian populations was evident from the Admixture analysis (Figure 2). As speculated, Punjabis (PJL), Gujaratis (GIH), and the ANI samples^23^ revealed a higher fraction of the ‘North Indian component’ (k1), in contrast, Tamils (STU), Telugus (ITU), and ASI samples^23^ appeared to possess a higher fraction of the ‘South Indian component’ (k2). To note, component k1 was found to be the highest among genomes of Kashmiri Pandits, Brahmins, Kshatriyas and Punjabis, while component k2 was higher among Paniyas, Palliyars, and Sri Lankan Veddas. Expectedly k3 that may be regarded as the North-East Asian component, was found to be the highest among the North-East Indians. Bengalis from Bangladesh, not only possess discernable ‘North Indian’ and ‘South Indian’ admixture components but have a significant fraction of component k3, which is potentially linked to their geographical origin and migration history. Component k4, that predominantly occurred among the Andamanese populations and was observed among most tribes from the Indian subcontinent except Bhils in discernible proportions. Finally component k5 was exclusively found among the Siddis. Since the Siddis are descendants of the Bantu tribe from Southeast Africa (Afro-Indians) and potentially arrived in India as a result of slave trade ~300-400 years ago^37^, their genomic exclusivity is not surprising. The geographic predominance of the various admixture components in the Indian subcontinent is shown in Figure 2 (inset).

**Figure 2.**
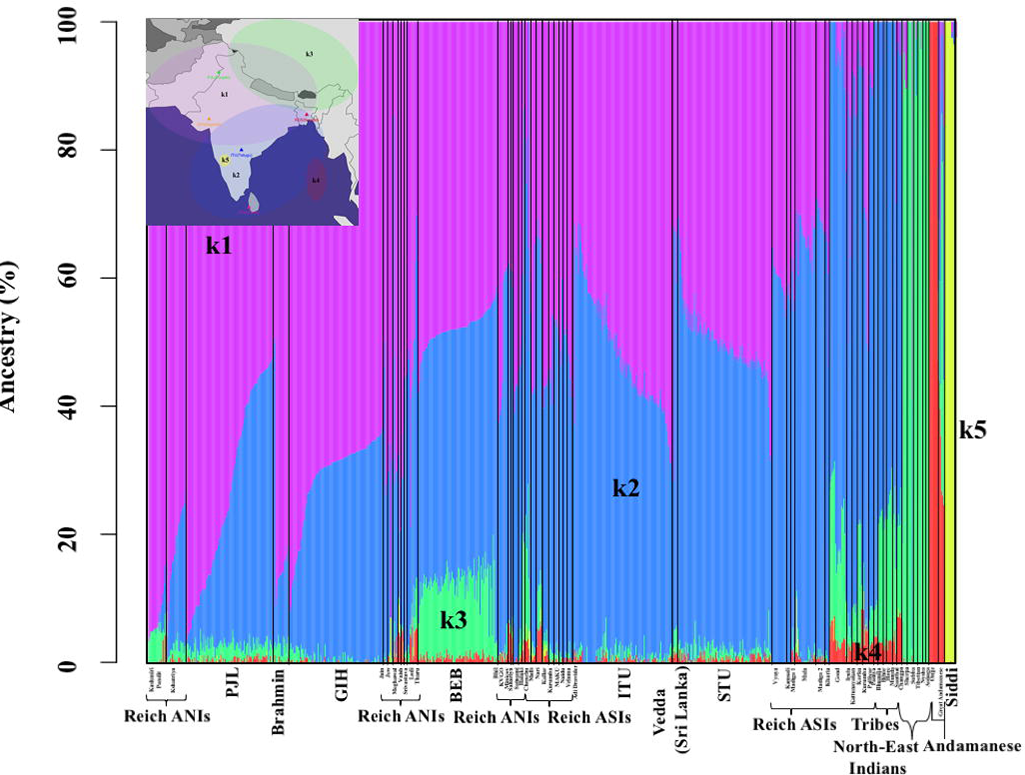
An Admixture plot showing the ancestry components of various South Asian populations. Percent ancestry is plotted on the Y axis. The ancestry of all 489 samples derived from 1000 Genomes project and 378 samples from Moorjani *et al.*, 2013^23^ was estimated using ADMIXTURE v1.3. To note k1, k2, k3, k4, and k5 represent putative ancestral North Indian, South Indian, North-East Asian, Andamanese, and African components respectively. **Inset:** A map of the Indian subcontinent showing the predominance of k1, k2, k3, k4, and k5 admixture components. The violet, blue, green, red, and yellow ellipses designate the putative circumference of k1, k2, k3, k4, and k5 components, respectively.

### Tracing the origin of South Asian populations

All biogeographical inferences were estimated using the GPS tool.^35^ The GPS assigned locations for all South Asian genomes evaluated are depicted in Figure 3. Most South Asian genomes were positioned in continental India, Lakshadweep islands and a few in Sri Lanka. Majority of Punjabi, PJL (>80%) and Gujarati, GIH (>70%) genomes were positioned in Northern and Western India, across the Indian states of Jammu and Kashmir, Uttarakhand, Uttar Pradesh, Rajasthan, and Gujarat. Strikingly the rest of the GIH genomes were positioned in Karnataka, Kerala, and Andhra Pradesh across the basin of the Krishna and Cauvery rivers. While the remaining PJL samples were positioned in Chhattisgarh (across the basin of the river Mahanadi) and in the Lakshadweep islands. Most Bengali, BEB genomes (~60%) were positioned in North India, spreading across the states of Uttarakhand, Uttar Pradesh and in Lakshadweep islands, while the rest were found scattered across South India with two individuals positioned in the Mahanadi basin and another in Sri Lanka. Expectedly most Tamil, STU and Telugu, ITU genomes were positioned in South India across the Indian states of Andhra Pradesh, Telangana, Tamil Nadu, Kerala, and Karnataka (across the Godavari, Krishna, Cauvery basins). While the remaining were positioned in North India across the Gangetic basin.

**Figure 3.**
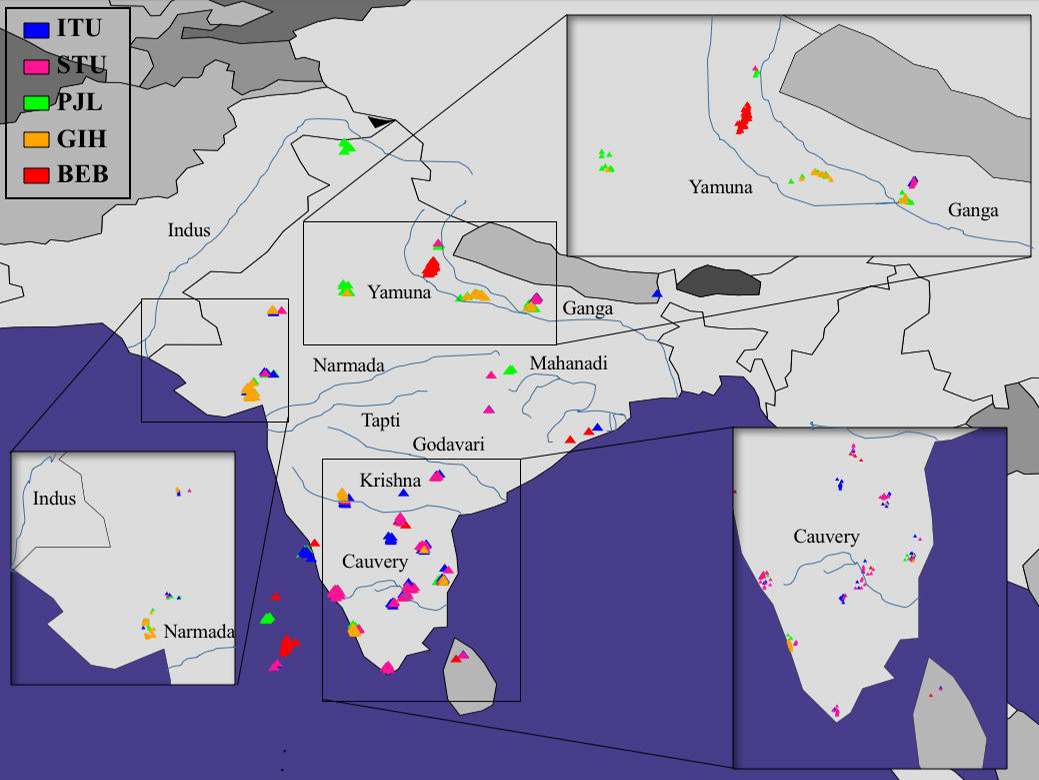
A map depicting the GPS assigned locations of the five South Asian populations from 1000 Genomes project. The red, blue, orange, green, and pink triangles depict BEB, ITU, GIH, PJL and STU populations, respectively. Prominent Indian rivers are depicted via blue lines (not to scale).

Interestingly, GPS seems to have positioned most South Asian genomes along trade routes that thrived in the ancient Mauryan empire, ~ 2300 YBP^38^ that unified the Indian subcontinent (Figure 4A) and are somewhat overlapping with the ancient purported migration routes of Proto-Dravidian and Indo-Aryan immigrants that eventually gave rise to ancestral South and North Indian populations, respectively (Figure 4B)^6,39^.

**Figure 4.**
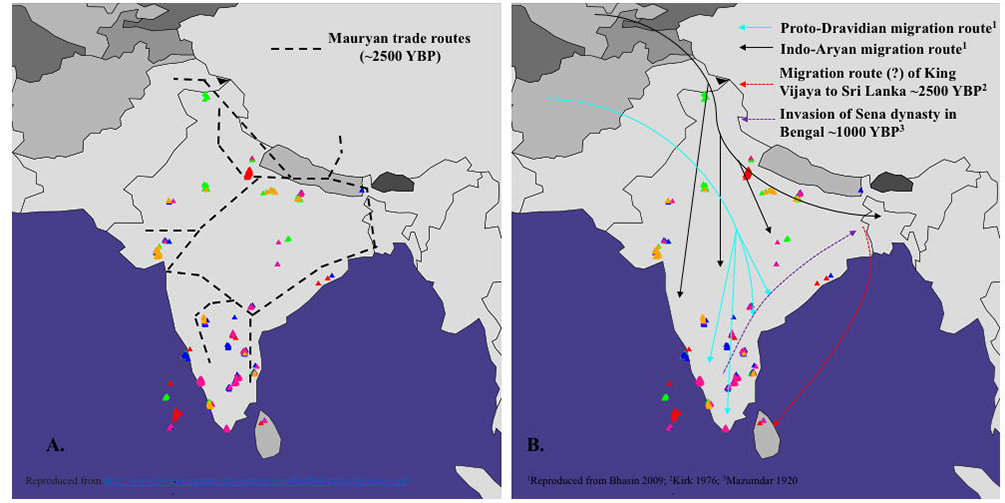
(**A**) A map depicting ancient Mauryan trade routes. The Mauryan trade routes are depicted in broken lines. **(B)** *A map showing ancient migration routes across Indian subcontinent*. The cyan and black lines represent the tentative migration routes of proto-Dravidians and Indo-Aryan immigrants respectively.^6,39^ The red and violet broken lines represent putative migration routes of groups from Bengal to South India and of people from Karnataka (South India) to Bengal, respectively.^46,48^

GPS assignment accuracy was determined for each individual to investigate how close they are positioned from their regional location (Supplementary Figure S1). At an average, more than 56% of all genomes were positioned within 600 km and only 10% were positioned more than 1500 km from their regional location (Supplementary Figure S1). These results illustrate reasonably strong genomic-geographic relationships and delineate the expected assignment error for the South Asian populations. The GPS prediction was most precise for ITU and STU samples where 87% and 96% samples respectively were positioned within 600 km of their native locations, Hyderabad and Colombo, respectively. Likewise, 79% GIH samples were positioned within 600 km of Ahmedabad, the capital of the Indian state of Gujarat. However, the GPS predictions were only moderately accurate for PJL and BEB samples. 19% Punjabis from Lahore were positioned within 600 km of it. Most PJL samples (69%) were positioned within 1200 kilometers of Lahore. Finally, 72% Bengalis from Bangladesh were positioned within 1500 kilometers of their native location, Dhaka, Bangladesh. We surmise that the moderate-low level accuracy for PJL and BEB samples could be attributed to the unavailability of Punjabi and Bengali genomes in the reference populations obtained from Moorjani *et al.*, 2013 ^23^.

## Discussion

Contemporary India and adjoining areas in South Asia are a panorama of extraordinary genetic, ethnic and linguistic diversity interwoven with a unique social structure. In the present study we have utilized the GPS tool to tease apart the complex biogeographical history of five populations from the Indian subcontinent. GPS positioned most South Asian genomes along the ancient migration routes of Proto-Dravidians and Indo-Aryans^39^ across the basins of the Ganges, Godavari, Krishna, and Cauvery rivers (Figure 3 and 4B). Interestingly, our GPS predictions also largely overlap with the sites where the skeletal remains of Central and South Indian communities (since 5000 B.C.), Megalithic builders and their Iron Age Collaterals (1300-500 B.C.) were uncovered.^39^ The most parsimonious explanation to account for this is that the admixture of ancestral North and South Indian populations primarily took place somewhere in Central and South Central India, along the basin of the Godavari and Krishna rivers that thus, may be regarded as the site where modern South Asian genomes likely originated.

GPS assigned most Southern Indian genomes, ITU and STU of Dravidic origin either to the Godavari river basin (Central-South India) or in the Southern Indian states of Kerala, Karnataka, and Tamil Nadu, while a few others were positioned in North-West and Central India (Figure 3). The strong assignment accuracy observed for these samples argues for the meaningfulness of this information and led us to hypothesize that this may be representative of ancient migration route(s) that eventually led to the peopling of the Southern part of the Indian subcontinent with Dravidian speaking populations. Our hypothesis is supported by archaeological, linguistic and genetic evidences that suggest that there were two prominent waves of immigrations to India. A majority of the Early Caucasoids were proto-Dravidian language speakers that migrated to India putatively ~ 6000 YBP.^6,7,39^ The migration of these groups from West Asia (Elam province) eastwards into India was a result of pastoral nomadism and the spread of agriculture.^40^ Further evidence for this migration comes from the Elamo-Dravidian languages, which may have originated in the Elam province (Zagros Mountains, South-Western Iran) but are now confined to South-Eastern India and some isolated groups in Pakistan and Northern India.^5,7,41^ It is noteworthy that the enigmatic script of the ancient Indus valley civilization (2600–1900 B.C.) that flourished in Pakistan and North-Western India, is also deemed to have similarities with ancient Dravidian languages.^42^ Finally the West Eurasian ancestry component has been estimated to be present in genomes from the Indian subcontinent prior to the purported Indo-Aryan migration, occurring ~3500 YBP, further suggesting that Eurasian populations migrated to India and admixed with the local populations before this time.^10^ Our findings are also favoured by previous theories of molecular anthropology that suggest an early migration of proto-Dravidian speaking Caucasoids to the Indian subcontinent, preceding the putative Indo-Aryan migration, followed by their gradual movement from North and West of the subcontinent to Central India, leading to initial settlements in the Godavari–Krishna rivers basin (Central-South India) and eventual migration further towards the Southern region of the Indian subcontinent, giving rise to the Dravidian South Indian populations.^39^

The Later Caucasoids or Aryans seemingly migrated to India ~3000-4000 YBP, initially localizing to the West of the river Indus, thereafter, settling along the banks of the river Ganges and its tributaries^12,39^ Towards the end of 6^th^ century B.C. the land of ‘Aryavarta’ or the country of Aryans was founded, bordered by Himalayas in the North and the Vindhya mountain range in Southern India. The predominant line of Aryan expansion at this time seemed to be along the South-West coast of India.^43^ After occupying Vidarbha in Central India, Aryans are proposed to have spread as far as the Godavari basin in Central-South India. ^39,43^ However, the incursions by the Aryans into the Southern Indian milieu were few and South India remained to be predominantly occupied by Dravidian populations. Our GPS assignments are in agreement with the putative route of the Late Caucasoid or Aryan migration and settlement in India. Accordingly most North Indian samples, Punjabis (PJL) and Gujaratis (GIH) that are of Eurasian origin are positioned in two major clusters: one along the banks of the Ganges and its tributaries (depicting the primary Aryan settlement) and another largely constituted by Gujaratis that appeared along the Godavari-Krishna-Cauvery rivers basin (Central–South India) where Aryans are proposed to have settled subsequently (Figure 3). Our results suggest that the foray of Aryan populations into the Godavari basin in Central-South India brought them in close proximity to Southern Indian populations.^12,39^ Historical evidences suggest that a few hundred years later, ~2300 YBP, the Mauryan empire was founded by king Chandragupta Maurya unifying the Indian subcontinent. The Mauryan empire was geographically extensive, in the North it was bordered by the Himalayas, in the West it extended to South-West Pakistan, South-East Iran and Afghanistan bordered by the Hindu Kush Mountains, in the East it spread to the Indian state of Assam and finally in Central-South India it extended to the Cauvery river basin^38^ It stands as one of the most significant periods in Indian history with thriving trade and agriculture^38^ that thus provided ample opportunities for admixture between the ancient North and South Indian populations. This is also congruent with the predicted timeline of admixture between ancestral North (ANI) and South (ASI) Indian populations that is said to have occurred ~1900-4200 YBP.^23^ The positioning of all South Asian genomes along ancient Mauryan trade routes in South Asia (Figure 4A) depicts a striking correlation between their population history and the economic and sociocultural dynamics of the Indian subcontinent ~2300 YBP.

GPS positioned Bangladeshi Bengalis (BEB) genomes predominantly in North India (Figure 3). This is consistent with previous studies suggesting that the gene-pool of BEB populations is more similar to mainland Indians than North-East Asian populations, even though it carries a discernible North-East Asian (Tibetan, Bhutanese, Nepalese and Chinese) ancestry component.^44,45^ The genomic similarity between Bengali, Telugu (ITU) and Tamil (STU) samples, as reflected in the multidimensional plot (Figure 1), is a relic of the ancient admixture between groups from Bengal and South India. This is supported by investigations suggesting a high degree of genetic closeness between Bengalis and the Sinhalese people of Sri Lanka.^46,47^ A prominent thread of historical evidence alluding to this is the purported migration of King Vijaya and his followers from Bengal to Sri Lanka and the subsequent influx of the ‘Bengali’ genomic component to South India (Figure 4B).^46^ Finally linguistic evidences also suggest that a major wave of migration originating in the Southern Indian state of Karnataka brought the Sena dynasty to Bengal thus promulgating admixture between Bengalis and South Indians (Figure 4B).^48^

Despite the five populations queried in this analysis being of South Asian lineage derived from individuals residing outside their native geographic locations in the Indian subcontinent, our assignment accuracy ranged from strong for Tamils (STU), Telugus (ITU) and Gujaratis (GIH) to moderate-low for Punjabis (PJL) and Bengali (BEB) genomes (the latter potentially due to lack of appropriate reference samples), reflecting an appreciable genomic-geographic correspondence and highlighting the robustness of the GPS algorithm for predicting ancestry. To the best of our knowledge this is the first study to have investigated the biogeographical origin of populations from the Indian subcontinent. Overall our findings provide key insight into the population history, economic and sociocultural forces that may have been instrumental in shaping the unique population structure of each of the five South Asian genomes evaluated.

## Titles and legends to figures

**Figure S1**. *Stacked bar plots representing the assignment accuracy of GPS algorithm*. Blue-violet, dark-blue, cornflower-blue, cadet-blue, cyan, and azure segments represent the positioning of samples from the 1000 Genomes project within 300 km, 600 km, 900 km, 1200 km, 1500 km and more than 1500 km, respectively from their corresponding native locations.

